# Experimental and computational analysis of calcium dynamics in 22q11.2 deletion model astrocytes

**DOI:** 10.1101/2021.09.16.460696

**Authors:** Ivan V. Maly, Wilma A. Hofmann, Mikhail V. Pletnikov

## Abstract

Intracellular calcium dynamics in spontaneously active cells such as neurons or astrocytes is an information-rich readout of the physiological state of the cell. Methods for deriving mechanistic information from biological time courses, as well as for algorithmically extracting cellular activity time courses from imaging data, have significantly advanced in recent years but been mostly applied to neuronal data. At the same time, the role for astrocytes, a type of glial brain cells, in cognition and psychiatric diseases remains poorly understood. Using calcium imaging, computer vision, and Bayesian kinetic inference, we analyze calcium dynamics in primary astrocytes derived from control or *Df1*/+ mice, a model of 22q11.2 deletion syndrome (DiGeorge syndrome). Inference of highest-likelihood molecular kinetic characteristics from the intracellular calcium time courses pinpoints a significant change in the activity of the sarcoendoplasmic reticulum calcium ATPase (SERCA). Applying a SERCA inhibitor to the control cells reproduces the differences detected in the deletion-bearing cells. Our work identifies for the first time the molecular changes driving the calcium kinetics in 22q11.2 deletion model astrocytes. We conclude that Bayesian kinetic inference is a useful tool for mechanistic dissection of a complex cellular phenotype, calcium dynamics, in glial cells. This method has the potential to facilitate formulation of specific hypotheses concerning the underlying molecular mechanisms, prioritization of experiments testing such hypotheses, and, in the future, individualized functional molecular diagnostics.

## INTRODUCTION

Intracellular dynamics of calcium ion concentrations is a paradigmatic self-organization phenomenon in eukaryotic cells (see, e.g., Chen and Zaikin, 2020; Maly, 2021). Modulation of calcium ion homeostasis in the cell is employed for perception, propagation, and processing of biological signals, for example, those originating from cell surface receptors. In many brain cell types, the baseline activity is not just a steady state of calcium fluxes between the cytosol and extracellular and intraorganellar spaces. In such spontaneously active cells, intracellular calcium concentrations undergo variations in the form of repeated elevations and regular or quasi-regular oscillations.

Calcium signaling in astrocytes has been linked to regulation of synaptic transmission (e.g., Arizono et al., 2020; Denizot et al., 2021; Ishibashi et al., 2019), neurodegeneration (Lattke et al., 2017; Li et al., 2018), neuroimmunological response (Ouali Alami et al., 2018), inflammation (Robb et al., 2020), and brain energy metabolism (Jha and Morrison, 2018; Maly et al., 2021; Takahashi, 2020). The contribution of astrocytes to the maintenance of adaptive levels of brain metabolism, in particular, breaks down in a number of major psychiatric disorders. Along with several other groups (e.g., Frintrop et al., 2019; Haroon et al., 2017; O’Leary and Mechawar, 2021; Rivera and Butt, 2019; Scofield and Kalivas, 2014; Verkhratsky et al., 2019; Verkhratsky et al., 2014), we have addressed a number of aspects of astrocyte involvement in the pathophysiology of schizophrenia and other severe psychiatric conditions (Jouroukhin et al., 2018; Jouroukhin et al., 2019; Reissner and Pletnikov, 2020; Shevelkin et al., 2020; Terrillion et al., 2017; Xia et al., 2016). In addition, we have previously addressed the molecular biology of the calcium system in other cells (Maly and Hofmann, 2016; Maly and Hofmann, 2018a; Maly and Hofmann, 2018b) and the fundamental properties of intracellular calcium kinetics from the quantitative dynamical systems perspective (Maly, 2021). Here, using live cell imaging and a new analytical pipeline that combines and adapts established computer vision and machine learning algorithms, we obtain some new results concerning the mechanism of altered spontaneous calcium dynamics in astrocytes affected by a genomic deletion that is a known genetic risk factor of schizophrenia.

The 22q11.2 microdeletion in humans is associated with a developmental syndrome affecting multiple organs. Also known as DiGeorge syndrome, it is among the highest-penetrance and highest-prevalence genetic factors of schizophrenia and other major psychiatric diseases (Bassett and Chow, 2008; Van et al., 2017). Reducing the dosage of approximately 60 genes, 22q11.2 deletion syndrome is a complex condition whose molecular mechanisms we are only beginning to unravel on the scale of genome-wide expression and proteomic interactions (Gokhale et al., 2019). The system-level studies point to neuronal mechanisms that affect development of synapses and maintenance of robust synaptic transmission, in agreement with the earlier investigations that identified alterations in neuronal calcium handling via modulation of expression of SERCA, the sarcoendoplasmic reticulum calcium ATPase (Devaraju et al., 2017; Earls et al., 2012). Although synaptic transmission in the brain depends on cooperation of neurons with astrocytes, whose processes envelop the neuronal synapses, little is known about how the 22q11.2 deletion affects glia. Considering the central role of astrocyte calcium signaling in the orchestration of the brain’s adaptation of its metabolism to the levels of synaptic activity, and the implication of these biochemical processes in the pathophysiology of schizophrenia and other major psychiatric diseases (see, e.g., Jha and Morrison, 2018; Maly et al., 2021; Takahashi, 2020), how astrocyte calcium signaling is affected in 22q11.2 deletion syndrome is an important outstanding question.

The methods of studying calcium signaling in brain cells, including astrocytes, now rely primarily on expression of engineered fluorescent calcium indicator proteins and imaging (see Khakh and McCarthy, 2015; Semyanov et al., 2020). Actuated by conformational transitions characteristic of their calmodulin moiety, these indicators have the required sensitivity to resolve intracellular calcium variations in the physiologically relevant concentration range (Chen et al., 2013). Sophisticated data processing algorithms are used in extracting time series information from video images. By and large, these data processing methods have been targeted at neurons, however, new algorithms are being proposed and tested that are designed for application to astrocytes. These specialized computational tools address the complexity of spatially propagated calcium signals in astrocytes when detecting calcium events in the video data (Bjornstad et al., 2021; Wang et al., 2017; Wang et al., 2019). Typically, however, astrocyte imaging data are processed manually or semi-manually by the investigator defining regions of interest (ROI) in the images, from which the fluorescent signal is then extracted using common ROI-based software routines (see, e.g., Eilert-Olsen et al., 2019; Semyanov et al., 2020). Although the semi-manual approach is widespread also in neuronal studies, multiple methods for computer vision analysis of neuronal data have been proposed (e.g., Apthorpe et al., 2016; Zhou et al., 2018). The ideal of parameter-free unsupervised extraction of neuronal calcium time series from imaging data was pursued in a recent development of an algorithm, MIN1PIPE, that is capable of handling typical neuronal video data after being supplied with only the spatial scale information (Lu et al., 2018). While development of astrocyte-specific image analysis methods is an important line of work, it appears feasible to attempt taking advantage of the capabilities of the advanced algorithms for neurons in investigations of astrocyte calcium.

Downstream analysis of calcium time series data has the potential to illuminate the kinetics of the molecular mechanisms that underlie observable calcium dynamics. Bayesian inference represents a powerful approach in this regard. Recently, a Bayesian approach to cell calcium kinetics has been fruitfully implemented in application to mammary epithelial cells (Yao et al., 2016). The classical Li-Rinzel kinetic model of intracellular calcium dynamics was used as a basis for the inference (Li and Rinzel, 1994). It was possible to infer the highest-likelihood values of kinetic parameters of channels and pumps responsible for calcium homeostasis and signal transduction in individual cells, and analyze the distributions of their values in cell populations. This method revealed existence of physiological heterogeneity in outwardly uniform cell cultures, likely representative of the widely acknowledged but still inadequately quantified individuality of cells in living tissues. A method for Bayesian inference of the kinetics that governs intracellular calcium has also been developed for neurons. The latter method relies on subordination of the neuronal calcium dynamics to action potentials. The corresponding algorithm, called CaBBI (Rahmati et al., 2016), is implemented as a module for a variational Bayesian package, VBA. VBA itself (Daunizeau et al., 2014) is a software tool of broad applicability, which can handle biological time series data of various kinds that arise in neuroscience. Both MIN1PIPE and CaBBI-VBA having been developed in the MATLAB programming environment, they appear to be ripe for computational integration in a pipeline capable of arriving at estimates of molecular kinetic parameters, starting from calcium imaging video data. In the present publication, we implement such a pipeline in application to studying 22q11.2 deletion astrocytes. This is achieved by modifying CaBBI to replace the neuron-specific action potential governing model with the Li-Rinzel model for spontaneous intracellular calcium dynamics, whose framework has been utilized extensively in applications to astrocytes (see Denizot et al., 2020; Manninen et al., 2018).

## METHODS

### Ethics statement

The animal procedures were performed in accordance with the current guidelines of the American Veterinary Medical Association and received prior approval of the institutional animal care and use committee of the State University of New York at Buffalo (approval number 2020-34).

### Cell culture and treatment

The *Df1*/+ mouse line, a transgenic model of the human 22q11.2 deletion (Lindsay et al., 1999; Paylor et al., 2001), was maintained on the C57BL/6J genetic background as previously described (Sumitomo et al., 2018). *Df1*/+ mice were kindly provided by Dr. Stanislav S. Zakharenko, St. Jude Children’s Research Hospital (Memphis, TN). Primary cultures were derived from cortices of neonate *Df1*/+ mice (bearing the hemizygous deletion) and their +/+ littermates (wild-type). The derivation of cultures was performed as previously described (Schildge et al., 2013). Specifically, the cortices were triturated, and the material digested with trypsin (Thermo Fisher, Waltham, MA) and seeded in flasks (Corning, Corning NY) coated with polylysine (Thermo Fisher, Waltham, MA).

Following one week in culture, during which the medium was changed, the cells were immunomagnetically purified using the Miltenyi Biotech (Auburn, CA) anti-ACSA2 astrocyte purification kit. Immunomagnetic purification was performed according to the kit manufacturer’s protocol, as also described previously (Batiuk et al., 2017; Jouroukhin et al., 2019). Purified cells were maintained in Sciencell (Carlsbad, CA) complete animal astrocyte medium.

After two weeks in culture, the cells were transduced with adeno associated viral vectors (AAV, multiplicity of infection 100,000) expressing the fluorescent fusion protein intracellular calcium indicator, GCaMP6m (Chen et al., 2013). The vectors were acquired from Signagen (Frederick, MD) and used in accordance with the supplier’s protocol. Specifically, the vector particles were added to the cell culture medium at the indicated concentration, and two days were allowed for expression before imaging.

For imaging, cells were grown in Labtek (Grand Rapids, MI) optical glass-bottom chambers coated with polylysine (Thermo Fisher, Waltham, MA). Prior to imaging, the medium was exchanged for Hanks balanced salt solution with calcium (1.26 mM), magnesium (0.89 mM), and D-glucose (5.6 mM, Thermo Fisher, Waltham, MA), and the culture was allowed to equilibrate for 40 min. In the experiments with thapsigargin (Thermo Fisher, Waltham, MA), the inhibitor was added to the medium at 200 pM on the microscope stage.

### Microscopy

Video recording was performed on a Leica DMi8 inverted fluorescence microscope equipped with an environmental control chamber for live cell imaging. The light-emitting diode excitation power was set to 1% and additionally attenuated by 85% using a neutral density filter to minimize light exposure. Images were acquired in a rapid time-lapse fashion (1 Hz) over 3 min with the excitation shutter closed between the frames, using Leica LAS X software and a monochrome cooled camera (Leica DFC9000GTC). In the thapsigargin experiments, a field of view was selected blindly and imaged before and 5 min after application of the inhibitor. In other experiments, the starting field of view was similarly selected and the culture imaged following a predetermined geometrical pattern around the first field, using the computerized stage control. Each field of view was recorded once, and a field diaphragm was used to limit excitation light exposure to the area being currently imaged.

### Image analysis

Imaging data were analyzed in MATLAB software (Mathsoft, Natick, MA). Individual active regions were automatically identified in the image sequences and fluorescence time courses extracted from them, using the MIN1PIPE package (Lu et al., 2018). MIN1PIPE is a computer vision algorithm for calcium indicator fluorescence signal extraction from video recordings of neural cells. It is designed to be essentially parameter-free in intended applications as long as the image scale is known. Version v3 alpha (https://github.com/JinghaoLu/MIN1PIPE/releases/tag/v3.0.0) was used with the default settings. The structural element size for the morphological opening operation was set according to the image scale (13 μm).

### Kinetic inference

#### Overview of the method

Kinetic parameters of the cell calcium system that are most likely, given the fluorescence time courses, were inferred using a suitably modified CaBBI module (Rahmati et al., 2016) for the VBA package (Daunizeau et al., 2014). VBA is an algorithm for analysis of biological time series and other types of data that implements a variational Bayesian approach (Friston et al., 2007) for these applications. CaBBI is a module for applying VBA to infer the cellular kinetics controlling the intracellular calcium concentration from fluorescent calcium indicator time series. VBA operates by describing a specific application with a generative model that consists of an evolution and observation functions, the first of which captures the intrinsic dynamics of the system in question and the second, the quantitative mapping of the intrinsic dynamics onto the measurement readout from the experiment. In a specific application, the VBA functionality is called by a MATLAB script specific to the problem, which also specifies the evolution and observation functions to be used. The CaBBI module supplies such a script and functions that are specific to calcium kinetics in neural cells.

The original CaBBI operates with kinetic models for action potential-driven calcium kinetics in neurons (Rahmati et al., 2016). To apply the same method to astrocytes, we replaced the action potential-based evolution models with the Li-Rinzel model for intracellular calcium kinetics (Li and Rinzel, 1994; see also, e.g., Nadkarni and Jung, 2007), which has been used widely in astrocyte calcium analysis (reviewed in Manninen et al., 2018). This model considers the exchange of calcium ions between the cytosolic and endoplasmic pools and incorporates the kinetics of SERCA, the inositol trisphosphate receptor (InsPR), and ligand-independent calcium leak out of the endoplasmic reticulum (ER). In the InsPR kinetics, intrinsic inactivation, potentiation by cytosolic calcium, and activation by inositol triphosphate (IP3) are captured. In addition, the Li-Rinzel model considers the production and degradation kinetics of IP3, the former being dependent on the cytosolic calcium concentration. We coded this model as an evolution function for use in the VBA-CaBBI environment. All parameters were held at their previously validated values (see Nadkarni and Jung, 2007), or varied around these values as prior means in the Bayesian inversion procedure. Specifically, the parameters that could be directly affected by the expression levels of the relevant proteins (extensive parameters) were varied: the rate constants of each of the calcium fluxes listed above, and of the production and degradation of IP3. Finally, to account for the properties of the calcium indicator used in our study, GCaMP6m, we set the value of the dissociation constant in the CaBBI observation function to 167 nM (Chen et al., 2013).

In sum, the Bayesian inference approach to calcium kinetics in astrocytes that is presented here is implemented by running CaBBI with a new evolution function in the MATLAB environment with the VBA toolbox installed. The new function encodes the kinetic equations described below.

#### Kinetic model

In the tradition referenced above (Manninen et al., 2018), our implementation of the kinetic model for spontaneous intracellular calcium variations in astrocytes follows Nadkarni and Jung’s (Nadkarni and Jung, 2007; Nadkarni et al., 2008) application of the Li-Rinzel model (Li and Rinzel, 1994). Specifically, the following differential equations are solved:

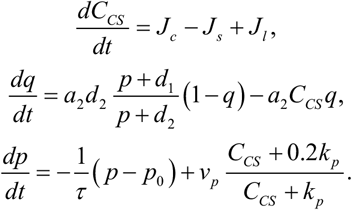

Here, *C_CS_* is the concentration of the calcium ion in the cytosol, *q* is the InsPR recovery variable, and *p* is the concentration of IP3. The model is closed by the following conservation relationship:

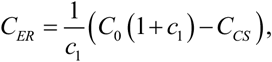

where *C*_0_ is the total concentration of calcium ion in cell and *c*1 is the volume ratio between ER and the cytosol. The fluxes through InsPR (*J_c_*), SERCA (*J_s_*), and the leak flux (*J_l_*) are defined as follows:

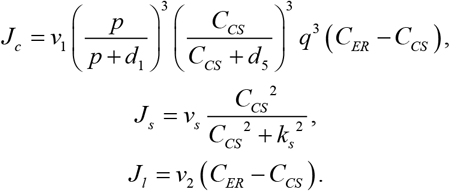

The values (Nadkarni and Jung, 2007; Nadkarni et al., 2008) of the parameters in the above expressions are given in Table 1.

**Table 1.**
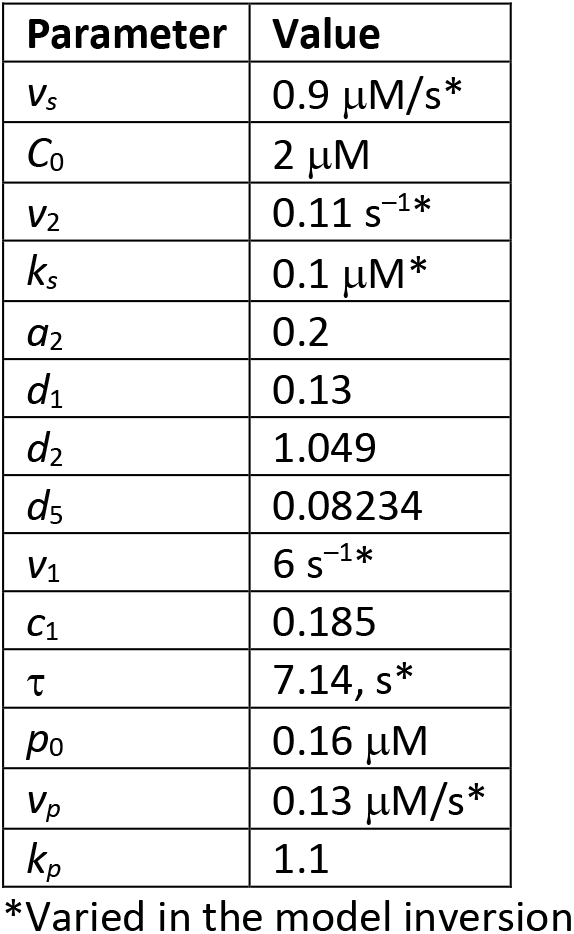
Model parameter values

#### Observation model

The cytosolic calcium concentration from the intracellular calcium kinetics model (*CCS*, above) is used to predict the calcium indicator fluorescence level. The CaBBI (Rahmati et al., 2016) approach to this task is encapsulated in the following observation function:

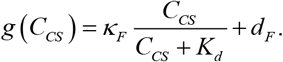

Here, *K_d_* is the dissociation constant of the indicator (set to 167 nM for our experimental conditions, see above), and *κ_F_*, *d_F_* are the scale and translation parameters that are determinable in the model inversion and account for the conditions specific to each observation (gain and background levels). The output of the observation function is evaluated directly against the recorded fluorescence in the model inversion.

#### Bayesian model inversion

In the VBA (Daunizeau et al., 2014) approach to approximate Bayesian inversion for biological time series models, hidden states *x* of the system are updated at each time point, *t*, using the deterministic evolution function *f* (kinetic model, above) and an assumption of additive noise:

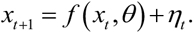

A similar approach is taken to the predictions of the observable, *y*, using the observation function *g*:

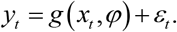

In these expressions, *θ* and *φ* are the parameters of the respective function, whereas *η* and *ε* are mean-zero Gaussian variables. Specifically, we have

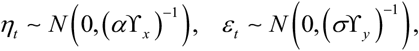

with Y being the corresponding inverse covariance matrices. In our case, *y* corresponds to the recorded fluorescence, while *x* is the set of dependent variables in the cell kinetic model (*C_CS_*, *q*, *p*). Appropriate accuracy of the kinetic model integration between time points *t* must be ensured, and this is achieved with the use of the “microtime” resolution setting as detailed below. Together, the evolution and observation models comprise the generative model, *m*.

The generative model inversion, given the data, is based on the variational Bayesian approach and Laplace approximation (Daunizeau et al., 2014). Given *m* and *y_t_*, we aim to infer the moments of the posterior distributions *p*(ν|*y_t_*, m) on ν = (*x_t_*, *φ, θ, α, β*). The conditional mean *μ* and covariance Σ are updated iteratively by optimizing *F*(*q*, *y_t_*) with respect to *q*(*ν*), the approximate posterior density. *F*(*q*, *y_t_*), termed free energy, constitutes the lower bound on the logarithm of the model evidence. It is computed as the difference of the log-evidence and the Kullback-Leibler divergence of the true and approximate posterior densities:

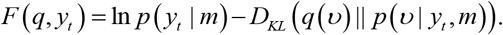

The method assumes that *q* can be represented as a product of the marginal posterior densities. The Laplace (Gaussian) approximation is used for each of these, except the precision parameters *α* and σ, which are modeled with gamma distributions. So, for the underlying kinetic parameters of interest, for example, we have

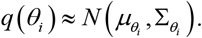

A version of the Gauss-Newton method is used for the iterative stepping, and a predefined limited number of subsequent observation time point data is used to update the hidden states at the given time point, instead of the full time series. Full computational details and references for the VBA method can be found in the cited work (Daunizeau et al., 2014).

#### Software versions and settings

MATLAB R2020b (Mathsoft, Natick MA) was used with VBA v1.9.2 (https://github.com/MBB-team/VBA-toolbox/releases/tag/v1.9.2) and CaBBI distributed as part of the same. Default parameter settings were used. Microtime resolution was selected as the method of integration of the underlying dynamical system (the Li-Rinzel model), whereby the integration step was 0.1 s, i.e., ten times smaller than the image acquisition interval.

#### Priors

In the VBA-CaBBI environment, the evolution parameter priors are Gaussian. The priors used are specified in Table 2. In accordance with the approach described above, the prior means of the kinetic parameters were equal to their values in the Li-Rinzel model. The other prior means and variances (Table 2) were either left at the values selected in the referenced version of CaBBI or—where the parameters such as the initial conditions were new—set to the correct order of magnitude according to the Li-Rinzel model (means) and CaBBI (analogous variances).

**Table 2.** Priors

| Parameter | Mean | Variance (on the exponent, where noted) |
| --- | --- | --- |
| $[\text{Ca}^{2+}]_{t=0}$, cytosolic | 0.1 $\mu\text{M}$ | 0.05 $\mu\text{M}^2$ |
| InsPR recovery variable at $t = 0$ | 0.5 | 0.2 |
| $[\text{IP3}]_{t=0}$ | 0.5 $\mu\text{M}$ | 0.2 $\mu\text{M}^2$ |
| SERCA $V_{\max}$ ($v_s$) | 0.9 $\mu\text{M}/\text{s}$ | 2 (exp) |
| InsPR $k$ ($v_1$) | 6 $\text{s}^{-1}$ | 2 (exp) |
| ER leak $k$ ($v_2$) | 0.11 $\text{s}^{-1}$ | 2 (exp) |
| IP3 $\tau$ (lifetime) | 7.14 s | 2 (exp) |
| IP3 synthesis $k$ ($v_p$) | 0.13 $\mu\text{M}/\text{s}$ | 2 (exp) |
| SERCA $K_m$ ($k_s$) | 0.1 $\mu\text{M}$ | 2 (exp) |

Non-negativity of the variable parameters was ensured through the transformation approach, as used in VBA and CaBBI. Specifically, the physical parameter was represented as a product of a constant equal to the parameter’s physical prior mean and an exponential factor whose exponent was the variable parameter in the computational inversion. Where this is the case, the logarithmic transformation is noted in Table 2. The dimensional prior mean value given in the table in these instances is the physical prior mean, whereas the corresponding computational inversion parameter had a zero prior mean and the dimensionless prior variance given in the table.

#### Convergence criteria

Individual GCaMP6 fluorescence traces constituted the datasets for independently run VBA inversions (one inversion per trace), resulting in inference of parameter sets characterizing the individual active regions. Each inversion was run for a maximum of 100 iterations using the default minimum increment on the free energy function (0.02). Runs returning a positive state noise precision hyperparameter were accepted. While not censored by fit of the data in the classical sense, it was informally verified that such runs resulted in an adequate fit (*R*^2^ > 90%). Informal analysis also showed that there was wide separation of the so-defined successful from unsuccessful runs in any dimension commonly used to evaluate data fit, and that the unsuccessful runs could be attributed to the comparative stiffness of the Li-Rinzel model and the relatively rapid stepping scheme selected for reasons of overall computational efficiency. The data (fluorescence traces) causing the rejected inversion runs were excluded from the analysis. In each experimental group, it was formally verified that the rejection rate did not exceed 5% (usually amounting to 0–3%).

#### Descriptive statistics and group comparisons

Posterior mean values of parameters estimated from fluorescence traces were treated in the frequentist framework as parameter sets characterizing the corresponding individual regions. This approach, previously employed in the mentioned Bayesian single-cell study (Yao et al., 2016), emulates standard handling of direct experimental cell measurements. Summary statistics was compiled across the active regions from the same experimental group. Group means and standard deviations calculated in this manner are presented for descriptive purposes. Statistical significance of group differences was evaluated using the Kolmogorov-Smirnov test on the distributions of the given parameter among the active regions of the experimental groups being compared. Specifically, active regions from wild-type cells were compared with active regions from deletion cells, while in the thapsigargin experiments, kinetics in the active regions before and after treatment was compared.

## RESULTS

Murine astrocytes expressing the fluorescent calcium indicator protein GCaMP6m that were derived from *Df1*/+ mice and their +/+ littermates (hereafter referred to as deletion-bearing and wild-type cells) exhibited robust spontaneous intracellular calcium dynamics in vitro. Identification of active calcium domains and extraction of their corresponding fluorescence time courses using the MIN1PIPE algorithm (Lu et al., 2018) showed that the individual active regions displayed a variety of dynamics. As seen in Fig. 1, the dynamics typically consisted of spikes that were widely separated in time or followed each other in periods of defined oscillations.

**Figure 1.**
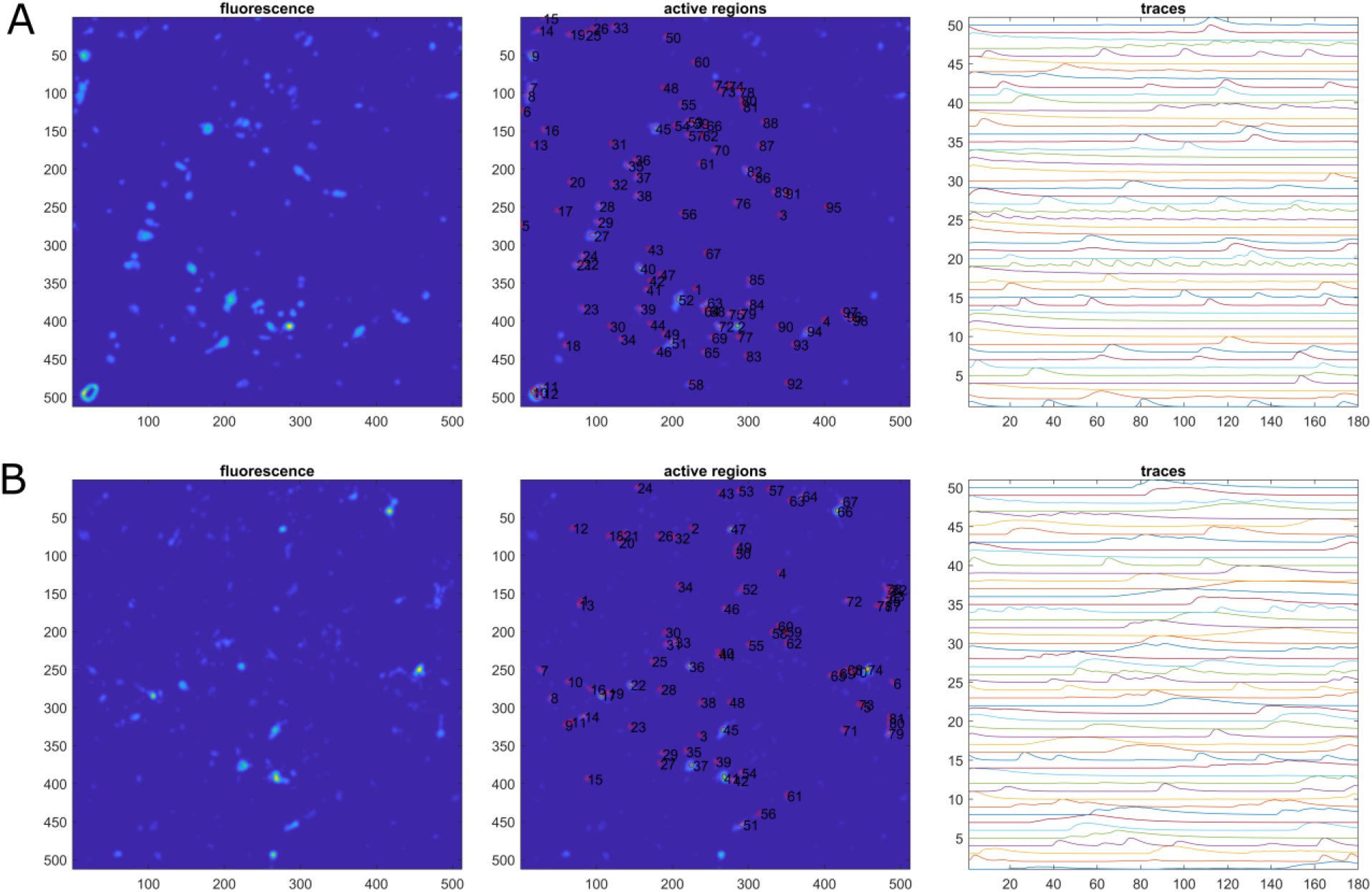
Calcium indicator fluorescence in wild-type and deletion-bearing astrocytes. A, wild-type cells, B, deletion-bearing cells. MIN1PIPE output modified for presentation. GCaMP6m fluorescence, active regions identified in the same fields, and time courses of fluorescence from the first 50 active regions are shown. Fluorescence images represent a maximum projection of the stack of time-lapse frames. The dimensions on the images are in pixels. Each image area is 670 by 670 μm. Fluorescence traces are normalized to the range. Time is in seconds, equal to frame numbers.

Bayesian inference of kinetic parameters of the Li-Rinzel model for astrocyte calcium dynamics (Li and Rinzel, 1994; Manninen et al., 2018) using the variational framework of VBA (Daunizeau et al., 2014) with a modified (see Methods) CaBBI calcium analysis module (Rahmati et al., 2016) allowed successful recapitulation of the recorded calcium indicator fluorescence traces and determination of the highest-likelihood parameter values of the underlying calcium kinetics (Fig. 2). The iterative approximate Bayesian inversion displayed consistent convergence, with the residuals displaying no significant temporal structure or autocorrelation (Fig. 3 C, G). The parameters of the observation function (*φ*, Fig. 3 H) were negatively correlated as expected for the scale and translation factors that are inferred from a certain time course of recorded fluorescence, given the time course of the intracellular calcium concentration. Correlation observed between certain pairs of kinetic parameters and initial conditions (*θ*, *x*_0_, see Fig. 3 H) reflected the intrinsic dependencies of intracellular calcium kinetics. SERCA and InsPR rate constants (*θ*_1_, *θ*_2_) as well as SERCA and ER leak rate constants (*θ*_1_, *θ*_3_, see Fig. 3 H) were positively correlated as expected for opposing fluxes near a steady state. The posterior correlation matrix (Fig. 3 H) further confirmed that the inferred SERCA *V*_max_ (*θ*_1_) and *K*_m_ (*θ*_6_) were negatively correlated, as expected for the maximum rate and Michaelis-Menten constant of an enzymatically-driven pump, given a certain value of the cytosol-to-ER calcium flux that can be inferred from the observation data.

**Figure 2.**
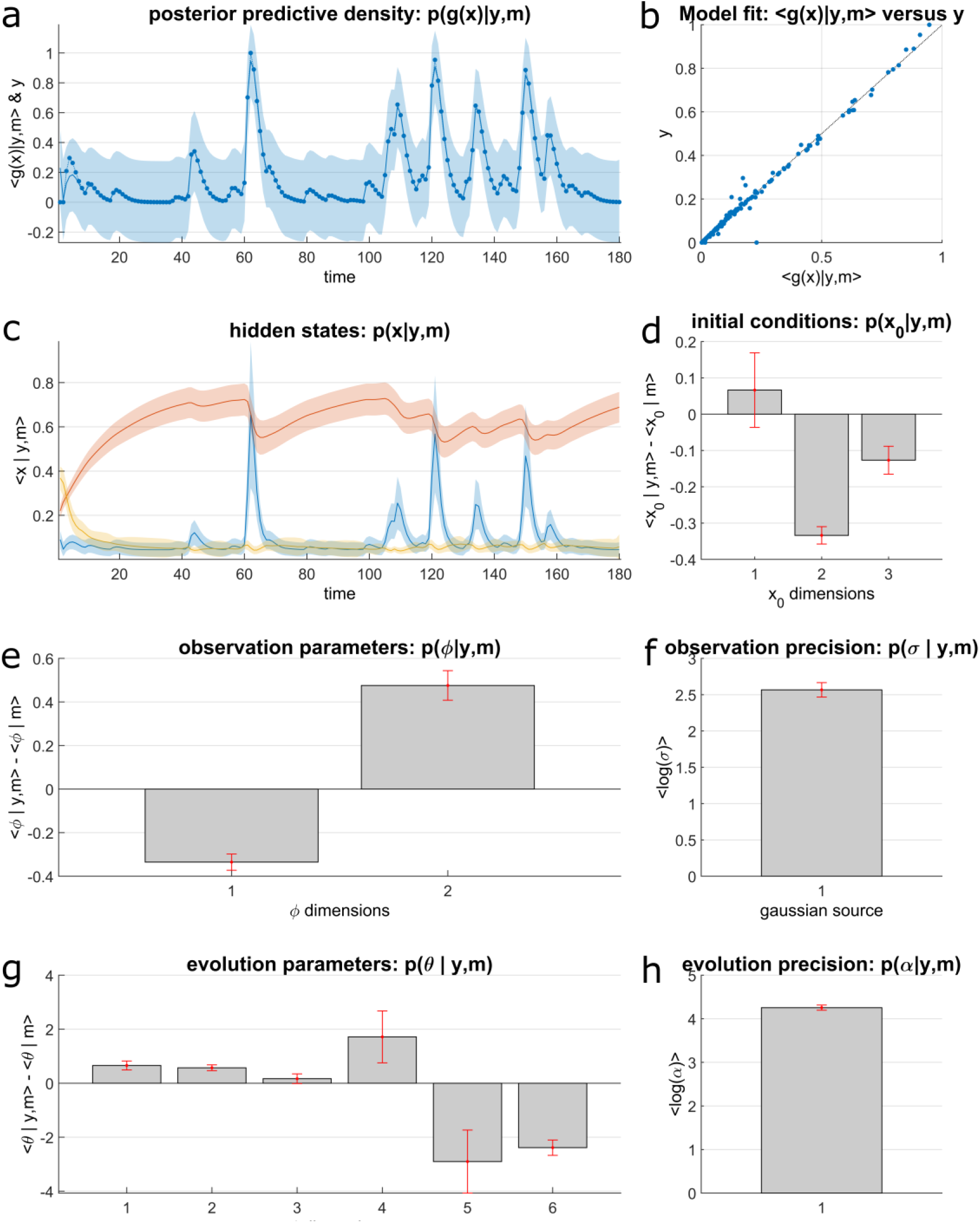
VBA inversion results for a wild-type active region. Observed (*y*) and predicted (*g*) indicator fluorescence (**a**, **b**) is dimensionless due to the normalizations employed. The hidden-states plots (**c**) are color-coded as follows: blue, cytosolic calcium concentration (μM), red, InsPR recovery variable, yellow, IP3 concentration (μM). Initial conditions (**d**) and evolution parameters (**g**) are plotted in the same order as they appear in Table 2. As explained in the text, the outputted posteriors on *θ* (the evolution parameters) are on the exponent in the factor applied to the prior mean to ensure positivity. The observation parameters (**e**) are the scale and shift parameters in the CaBBI observation function.

**Figure 3.**
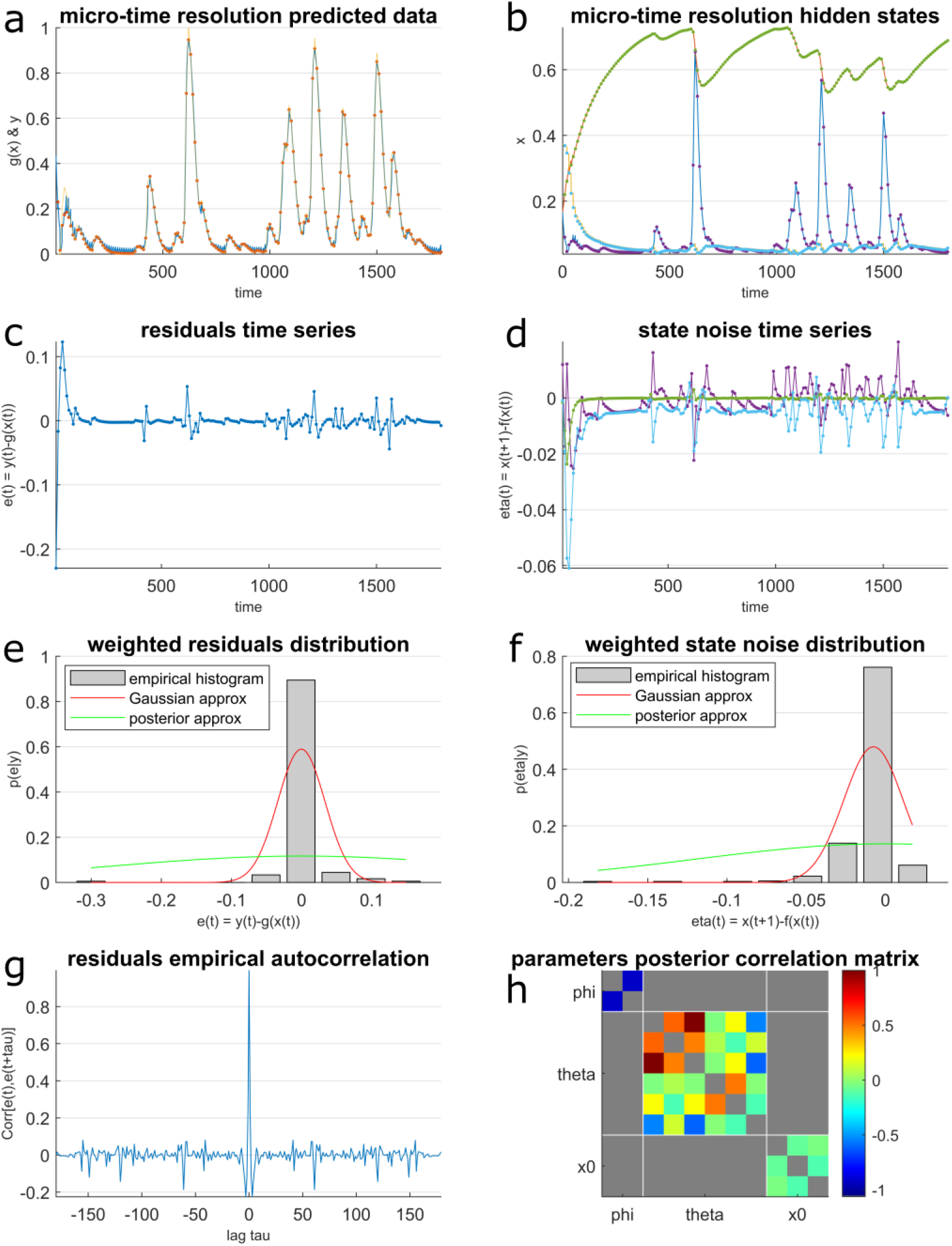
VBA diagnostics results for the same region. Hidden states and state noise plots (**b**, **d**): purple dots, cytosolic calcium concentration (μM), green, InsPR recovery variable, blue, IP3 concentration (μM). In the correlation matrix (**h**), *φ* are the scale and translation parameters of the observation function. As before, the evolution parameters (*θ*) and initial conditions (*x*_0_) are in the same order as they appear in Table 2. See the text for a discussion of the physics and biochemistry behind the parameter correlations.

Individual active regions were distinguished by their sets of inferred kinetic parameters. Statistics of the posterior means associated with the individual active regions were derived (Table 3). The complete dataset comprised recordings made on independently derived cultures from 7 deletion-bearing and 5 littermate background-strain mice. Overall, the inferred kinetic parameters displayed similar distributions in the populations of wild-type and deletion cells (Fig. 4). At the same time, a leftward (to the smaller values) shift of the distributions of the maximum rate of the ER calcium pump (SERCA *V*_max_) and rate constant of the ligand-independent flux of calcium out of the ER (ER leak *k*) could be seen, which was comparatively pronounced (Fig. 4 A, C). These two parameter shifts were associated with especially low *p* values in the Kolmogorov-Smirnov test that evaluates the separation of distribution functions (*p* < 10^−20^). It should be noted that the less pronounced shifts of the other parameter distributions also achieved statistical significance (*p* < 10^−2^). Assessing the significance alongside the relative change in the parameters (Fig. 5) showed that the parameter differences were clustered, with most falling into the group of small effect sizes and larger *p* values. SERCA *V*_max_ and ER leak *k* constituted the second cluster at a notably higher absolute effect size and significance. Evidently, while the inferred kinetic perturbation in the deletion-bearing cells was complex, these two parameters were especially notably changed.

**Table 3.** Summary statistics of individual posterior means

| Parameter | Untreated wild type |  | Deletion |  | Thapsigargin |  |
| --- | --- | --- | --- | --- | --- | --- |
|  | average | SE | average | SE | average | SE |
| SERCA $V_{\max}$, $\mu\text{M}/\text{s}$ | 1.57 | 0.02 | 1.34 | 0.01 | 1.26 | 0.02 |
| InsPR $k$, $\text{s}^{-1}$ | 13.9 | 0.1 | 13.3 | 0.1 | 11.9 | 0.3 |
| ER leak $k$, $\text{s}^{-1}$ | 0.108 | 0.001 | 0.090 | 0.001 | 0.084 | 0.002 |
| IP3 $\tau$ (lifetime), s | 34.0 | 0.2 | 33.5 | 0.1 | 32.1 | 0.4 |
| IP3 synthesis $k$, $\mu\text{M}/\text{s}$ | 0.00782 | 0.00002 | 0.00796 | 0.00002 | 0.00775 | 0.00006 |
| SERCA $K_m$, $\mu\text{M}$ | 0.0118 | 0.0001 | 0.0127 | 0.0001 | 0.0123 | 0.0002 |

**Figure 4.**
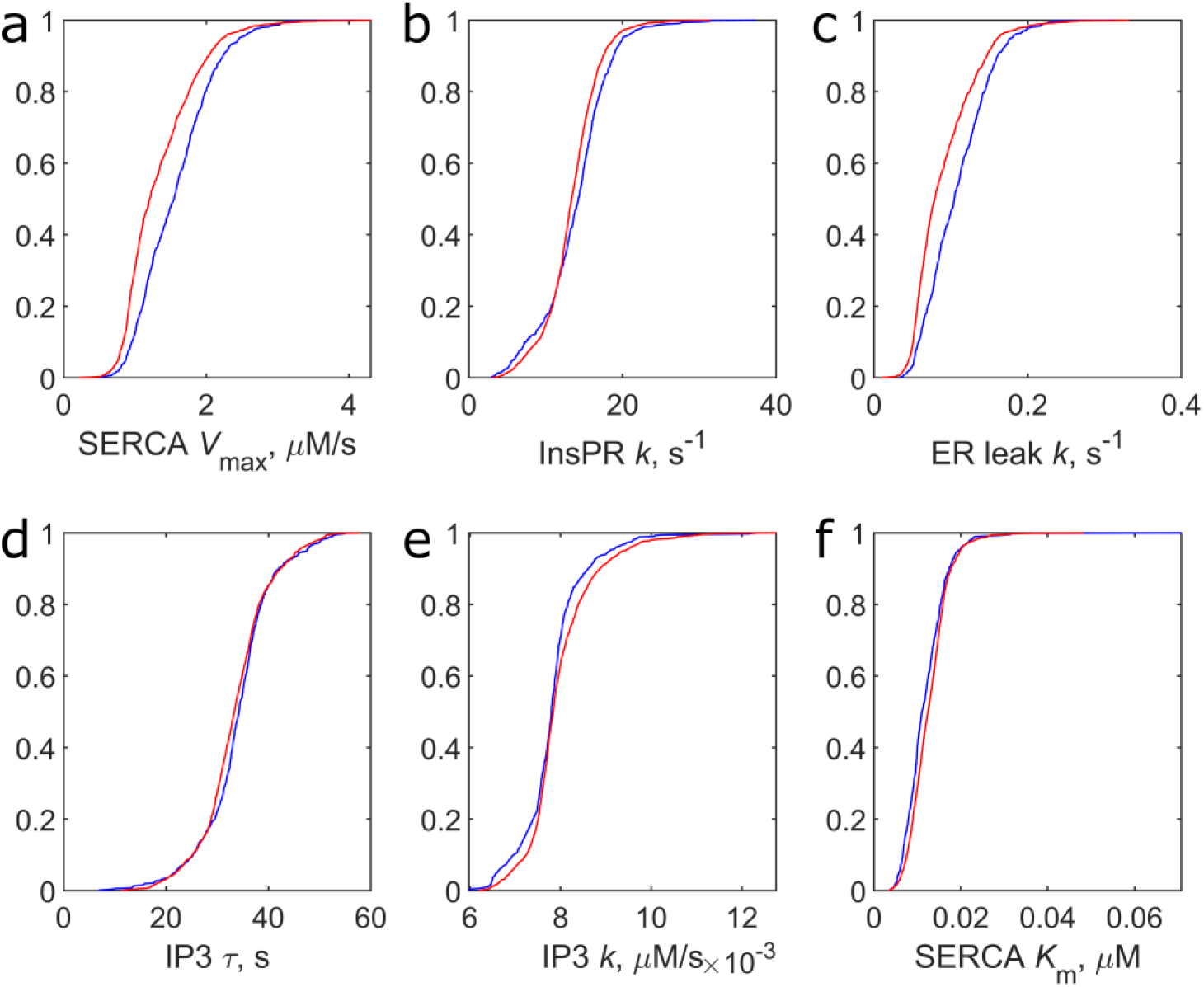
Distribution functions of calcium kinetics parameters in wild-type and deletion-bearing astrocytes. Blue, wild type, red, deletion.

**Figure 5.**
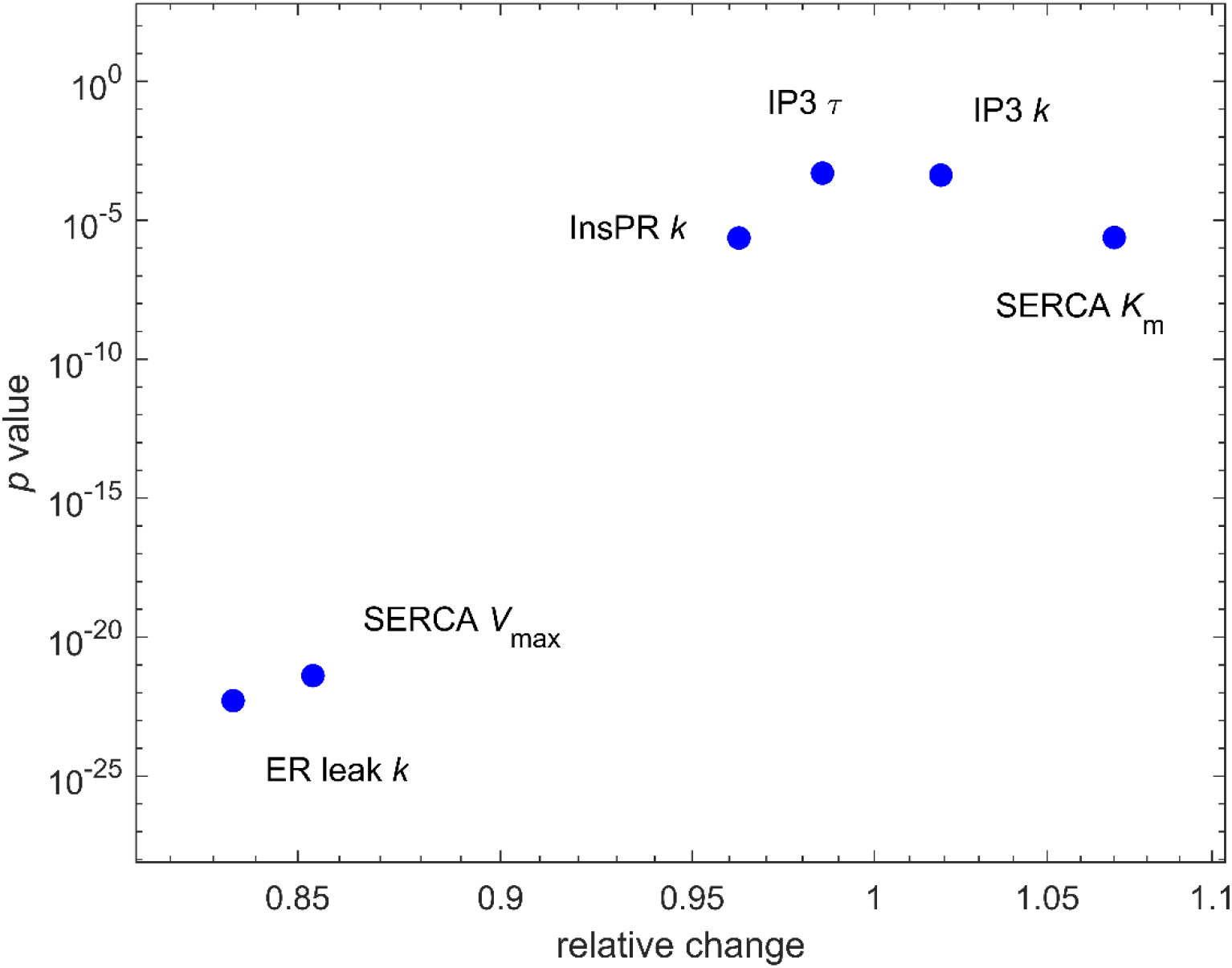
Shift of kinetic parameters in deletion-bearing cells. Ratios of population averages on individual posterior means and significance in the Kolmogorov-Smirnov test are plotted.

To test the validity of the kinetic inference pinpointing the effect on SERCA experimentally, we applied a well-characterized and widely used SERCA inhibitor, thapsigargin (reviewed in Andersen et al., 2015), to wild-type cells. The dataset collected as described in Methods was based on recordings from 127 calcium activity regions detected before application of thapsigargin and 227 regions in the same fields that were detected after application of the inhibitor in three separate experiments. Juxtaposition of the intracellular calcium indicator data from the cells before and after treatment (200 pM for 5 min) again revealed a wide variety of responses and individual dynamics (Fig. 6). Fluorescence traces from the thapsigargin-treated cells were similarly amenable to Bayesian inversion, and statistics of the inferred parameter values before and after the treatment were derived (Table 3). Their distributions (Fig. 7) showed similar forms before and after the inhibitor treatment, except for SERCA *V*_max_ and ER leak *k*, which were markedly shifted toward smaller values (Fig. 7 A, C). In the significance-effect size plot (Fig. 8), the cluster to which most of the parameters belonged had a similar composition to the one observed in the deletion vs. wild-type comparison (cf. Fig. 4). This group of parameters displayed a somewhat larger variation of effect sizes, but their statistical significance was lower (*p* > 10^−2^), and could not withstand a Bonferroni correction for multiple comparisons. At the same time, SERCA *V*_max_ and ER leak *k* again constituted a separate group at higher absolute effect sizes and significance (*p* < 10^−6^, see Fig. 8).

**Figure 6.**
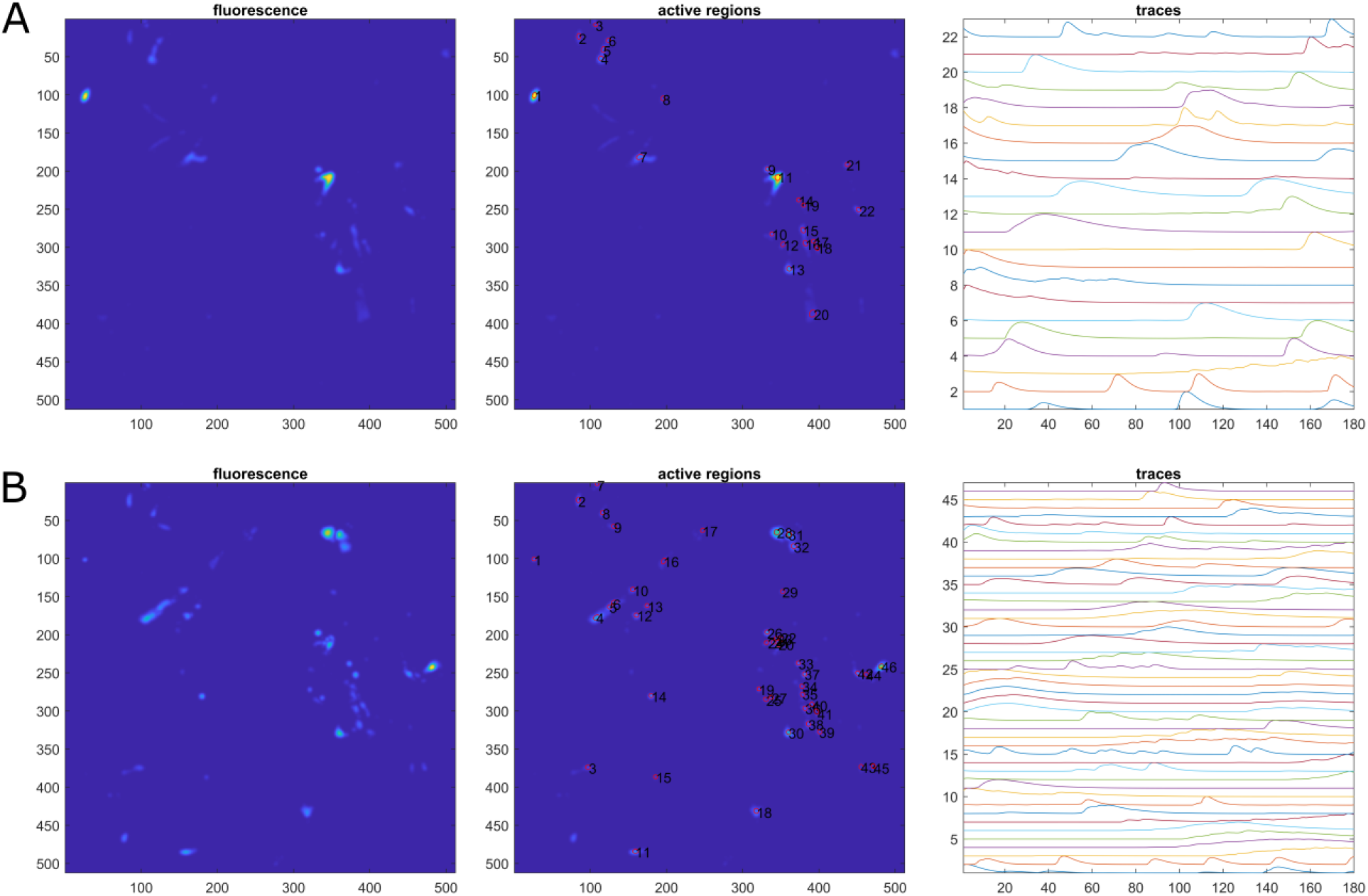
Calcium indicator fluorescence in wild-type astrocytes before and after treatments with thapsigargin. A, before application of thapsigargin, B, same field 5 min after application of 200 pM thapsigargin. Plotting as in Fig. 1.

**Figure 7.**
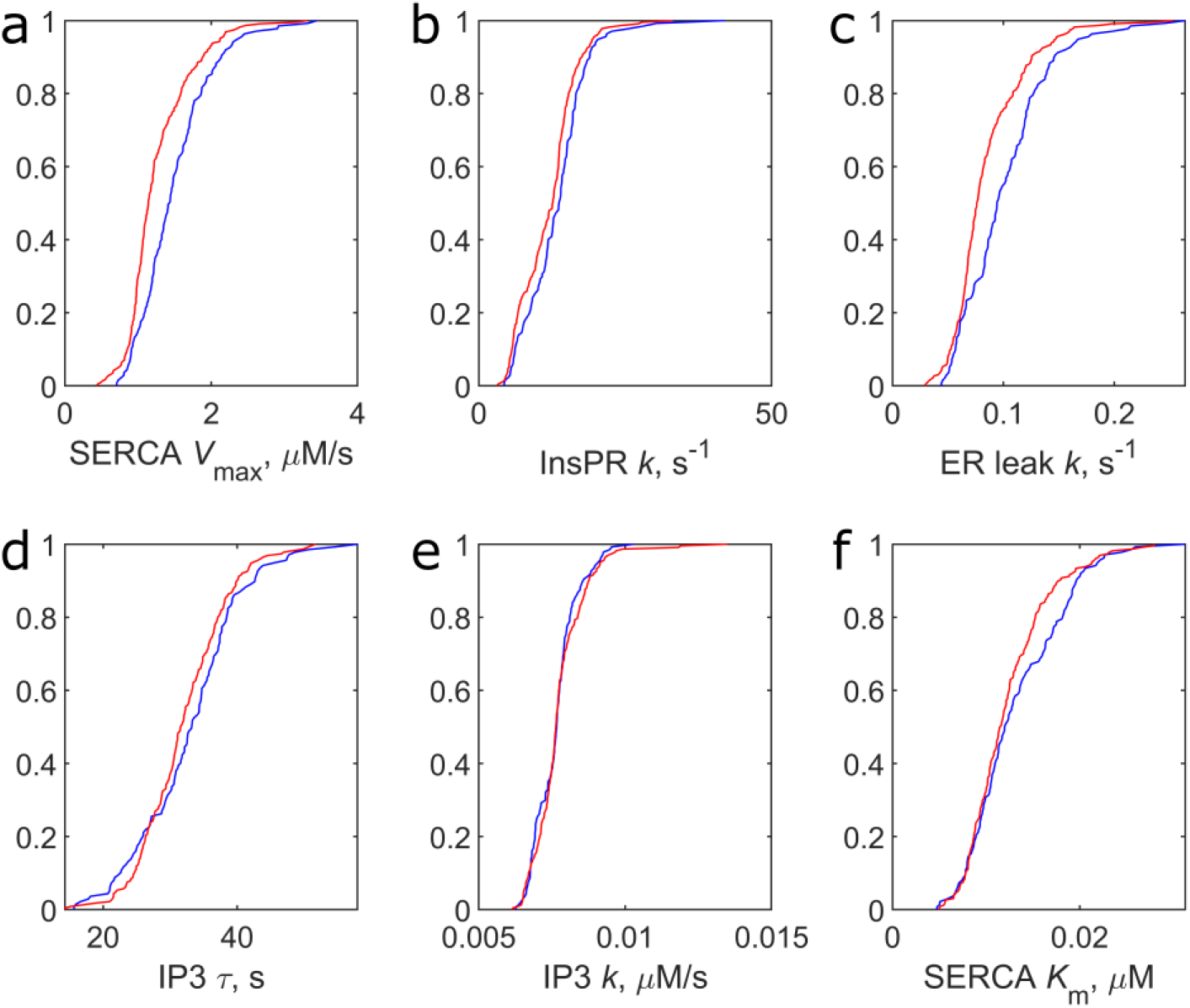
Distribution functions of calcium kinetics parameters in wild-type astrocytes before and after treatment with thapsigargin. Blue, before, red, 5 min after application at 200 pM.

**Figure 8.**
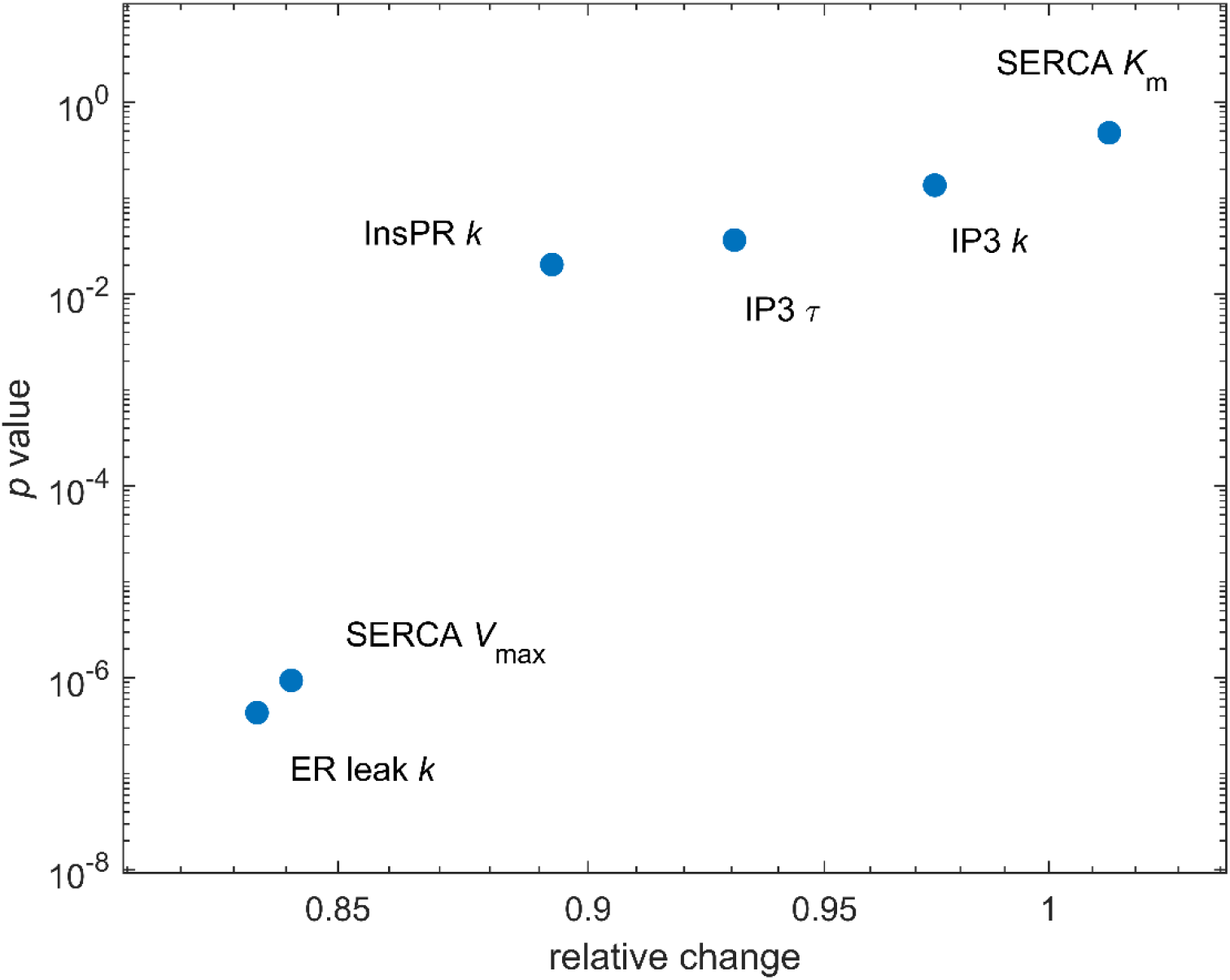
Thapsigargin-induced change in kinetic parameters

## DISCUSSION

The Bayesian kinetic inference approach proved capable of identifying salient molecular-level perturbations of intracellular calcium dynamics in a murine transgenic model of 22q11.2 deletion syndrome. Despite considerable cell-to-cell variability in spontaneous, continuous calcium dynamics, we have identified SERCA *V*_max_ and ER leak *k* as the parameters the values of which have been significantly decreased in the populations of deletion model astrocytes. Experimental tests using thapsigargin confirmed that the inferential analysis correctly identifies the altered molecular-kinetic parameters. Given the central role of the astrocyte calcium signaling system in astrocyte physiology and pathophysiology (see, e.g., Jha and Morrison, 2018; Maly et al., 2021; Takahashi, 2020) and the prominence of 22q11.2 deletion in mental illness (Bassett and Chow, 2008; Van et al., 2017), pinpointing the kinetic changes of the astrocyte calcium system in a murine model of this genetic syndrome is a new and significant finding that demonstrates the promise of the chosen analytical approach.

Dependencies between the kinetic parameters that are intrinsic to the intracellular calcium system were correctly reflected in the posterior correlation matrix, as described under Results. Since the two parameters that were found to be significantly altered in deletion-bearing cells are among the correlated parameter pairs, the nature of their mutual dependence merits an additional discussion. Diagnostics of our Bayesian inversion using VBA warns us that variations of SERCA *V*_max_ and ER leak *k* could not be inferred separately, given the data and the model (see Fig. 3 H). The high degree of inseparability implied by the correlation reflects, as already noted, the construction of the intracellular calcium system and the principles of its dynamics. Indeed, despite the considerable information embodied by the time courses recorded in our experiments, the cells maintain homeostasis of their calcium system through the variations, and the underlying dynamics remains near a steady state. Opposing fluxes near the steady state are bound to be correlated. Although this can be achieved by the relevant concentrations changing in concert, a change in the kinetic constant of one flux can also be compensated by a change in its opposite number.

Another reason for co-regulation of the two parameters in question may lie beyond the Li-Rinzel model. In this model, like in other frequently applied cell-scale calcium models (Li and Rinzel, 1994; Manninen et al., 2018), the opposing fluxes affected by SERCA *V*_max_ and ER leak *k* are formally independent, and inseparability arises only from the near-steady-state condition. In reality, however, operation of SERCA is bidirectional and the reverse flux of calcium through this pump is a component of the ER leak current (Camello et al., 2002). Indeed, suppression of ER leak is a known property of thapsigargin, which, naturally, inhibits both forward and reverse rate of SERCA (Camello et al., 2002). In view of this, one should expect co-regulation of the two fluxes whenever SERCA operation is affected as we find in our experimental model. In the present case, the fact that the experimental application of thapsigargin is capable of recapitulating the relative magnitude of the changes in the two parameters that is associated with the genomic deletion (Table 2, cf. Fig. 5 and 8) is compatible with a notion that the leak suppression by the deletion is effected via the bidirectional operation of SERCA rather than via any other possible mechanism of the ER calcium leak current (see Camello et al., 2002).

Two remarks are necessary in connection with our discovery of the effect on SERCA and use of thapsigargin to validate the discovery method. Firstly, SERCA is not among the genes whose dosage is affected by the deletion; see the 22q11.2 and transgenic model gene lists in, e.g., Earls et al. (2012). The effect detected here, therefore, is indirect and possibly mediated by the expression regulatory pathway previously characterized in neurons (Earls et al., 2012). Secondly, the concentration of thapsigargin employed in our confirmatory tests is dictated by the context of the subtle modulation rather than a shutdown of spontaneous calcium activity in the comparison of deletion-bearing and wild type cells. This concentration, therefore, is much lower than that employed in many cell biological experiments. Effectiveness of thapsigargin in the picomolar range, however, is well documented on the molecular level (Davidson and Varhol, 1995).

Our findings are consistent with prior results obtained in 22q11.2 deletion model neurons, in which altered calcium signaling in modified paired pulse facilitation (Gokhale et al., 2019). *Mrpl40* haploinsufficiency and age-dependent microRNA mechanisms were found to impact calcium dynamics, with both mitochondrial buffering in presynaptic terminals and SERCA expression being altered (Earls et al., 2010; Earls et al., 2012). However, while a plausible explanation has been offered to the detected SERCA overexpression being accompanied by prolonged calcium transients in the terminals, it has also been noted that this direction of change is opposite to how the pump usually operates by limiting the duration of calcium transients (Earls et al., 2010). Although the molecular pathway affecting SERCA expression in 22q11.2 neurons could operate in astrocytes as well, a different outcome may be observed due to cell type specific regulation. Following clarification of the astrocytic and neuronal mechanisms, future studies may be able to address the question how calcium dynamics alterations in the two cell types combine to affect synaptic transmission in 22q11.2 deletion syndrome.

While our focus is on biological applications and results, it is worth comparing our new computational pipeline with that in the cited recent study on a mammary epithelial cell line (Yao et al., 2016). The two approaches have substantially similar overall functionality in Bayesian inference of molecular cell-biological kinetics from calcium imaging data. The cited work did not employ direct detection of intracellular calcium activity in the image data, using instead image segmentation based on detection of nuclei with a DNA intercalating dye. This has a potential advantage of referencing the results to specific cells in the experiment, and to the overall number of cells, including inactive ones. This possibility is of less relevance in application to astrocytes with their characteristic independently active calcium domains, whose individual activity is believed to determine the local interaction with specific neighboring neurons and other cells in the brain (Khakh and McCarthy, 2015; Shigetomi et al., 2016). Further, the implementation developed for epithelial cells (Yao et al., 2016) similarly used the Li-Rinzel model as a basis for the kinetic analysis, and similarly employed approximate Bayesian computation, owing to the complexity of the nonlinear dynamics. The computation was, however, Monte Carlo-based rather than variational. The Monte Carlo approach, as demonstrated successfully in the cited work, is especially powerful in identification of multiple maxima of likelihood, whereas the variational algorithm may be better suited to the problem of defining continuous and subtle variability among the cells, as reported here. Finally, it may be worth mentioning that the cited approach employed a Python package for Bayesian inversion, while our pipeline is integrated in the MATLAB environment. Overall, while the differences contrasted here are not expected to be critical in all applications, our approach and that of Yao et al. (2016) offer somewhat different capabilities that can be tuned to the cell types being studied and computational resources of the laboratory. Together, they demonstrate that Bayesian analysis of cell calcium kinetics is a general method that can be implemented according to the needs of specific future studies.

The new application of the Bayesian kinetic inference method shows the promise of this method in analyzing the molecular underpinnings of complex genetic conditions. The combination of computational inference with the quality of imaging data and accessibility to experimental perturbations that are afforded by studying neural cells in vitro has the power to unlock the complexity that is the mark of genetic causation of mental illness. Further application of this method can allow rapid, rigorous formulation of specific hypotheses concerning the underlying molecular mechanisms and assigning rank priority to such hypotheses for experimental testing. The efficiency of the kinetic inference method has an obvious advantage over the traditional approach. Instead of formulating mechanistic hypotheses in the order not directly dictated by the data, and instead of testing them by comparing descriptive characteristics of calcium dynamics observed in the tests, the inferential approach starts with a pair of datasets (e.g., deletion and wild type) and ranks the likelihood of specific molecular-level differences that may underlie the data. These molecular-level possibilities can then be tested experimentally in the order of their objective likelihood. The example presented here demonstrates that the objective ranking process can even reduce a potentially very multifactorial system to one or two molecular-level effects, which can then be rapidly tested experimentally. Given its ability to detect kinetic differences in individual cell cultures, furthermore, application of this method to patient-derived cells can be an avenue to functional molecular diagnostics as part of future individualized medicine approaches. At the same time, our demonstration of kinetic inference from time series data supplied by a computer vision algorithm capable of handling noisy in vivo images establishes an analytical pipeline that appears ready for in vivo applications. Systematic determination of molecular kinetic parameters underlying calcium dynamics in spontaneously active cells such as astrocytes under different physiological, genetic, and pathophysiological conditions will constitute a new level of quantitative understanding of this paradigmatic self-organization phenomenon in the cell.

## ACKNOWLEDGMENTS

This work was supported in part by the NCI of the National Institutes of Health under award number 254 R21CA220155 to Wilma A. Hofmann and NIMH of the National Institutes of Health under award number R01MH083728 to Mikhail V. Pletnikov.

## Notes

### Competing Interest Statement

The authors have declared no competing interest.

### Summary of Updates

The Methods section has been expanded.

